# *In vitro* evaluation of therapeutic antibodies against a SARS-CoV-2 Omicron B.1.1.529 isolate

**DOI:** 10.1101/2022.01.01.474639

**Authors:** Franck Touret, Cécile Baronti, Hawa Sophia Bouzidi, Xavier de Lamballerie

**Affiliations:** Unité des Virus Émergents (UVE: Aix-Marseille University - IRD 190 - Inserm 1207), Marseille, France

**Keywords:** SARS-CoV-2, Omicron, therapeutic antibody, variant

## Abstract

The emergence and rapid spread of the Omicron variant of SARS-CoV-2, which has more than 30 substitutions in the spike glycoprotein, compromises the efficacy of currently available vaccines and therapeutic antibodies. Using a clinical strain of the Omicron variant, we analyzed the neutralizing power of eight currently used monoclonal antibodies compared to the ancestral B.1 BavPat1 D614G strain. We observed that six of these antibodies have lost their ability to neutralize the Omicron variant. Of the antibodies still having neutralizing activity, Sotrovimab/Vir-7831 shows the smallest reduction in activity, with a factor change of 3.1. Cilgavimab/AZD1061 alone shows a reduction in efficacy of 15.8, resulting in a significant loss of activity for the Evusheld cocktail (42.6 fold reduction) in which the other antibody, Tixagevimab, does not retain significant activity against Omicron. Our results suggest that the clinical efficacy of the initially proposed doses should be rapidly evaluated and the possible need to modify doses or propose combination therapies should be considered.

## Short communication

Since the emergence of the SARS-CoV-2 coronavirus in China in late 2019, the virus has spread worldwide, causing a major pandemic. The epidemic spread has been supported by the appearance of variants that combine increased transmissibility and antigenic escape to varying degrees. At the time of writing, we are witnessing the rapid replacement of the delta variant by a new variant, Omicron, which has a higher transmission capacity than all the previous variants, but also has substantial antigenic changes. Omicron has been first characterised in South Africa (B.1.1.529 lineage,”Weekly epidemiological update on COVID-19 - 21 December 2021,” n.d.) and exhibits the highest number of genomic mutations reported so far, especially in the spike glycoprotein where over 30 substitutions are present (Kumar et al., 2021). Such changes in the most important antigen of the virus, against which the neutralising humoral response is built, have the potential to significantly reduce the efficacy of both vaccines and therapeutic antibodies currently in clinical use (Malani et al., 2021; Taylor et al., 2021), as most of them were designed from the spike protein of the original SARS-CoV-2 strain (Baum et al., 2020; Cathcart et al., 2021; Jones et al., 2021; Kim et al., 2021).

In the current study, we tested the neutralising activity of a panel of COVID-19 therapeutic antibodies against a clinical strain of the Omicron variant. The ancestral D614G BavPat1 European strain (B.1 lineage) was used as a reference to calculate the fold change between the EC_50_s determined for each virus. To do this, we applied a standardised methodology for evaluating antiviral compounds against RNA viruses, based on RNA yield reduction (Delang et al., 2016; Kaptein et al., 2021; Touret et al., 2019), which has been recently applied to SARS-CoV-2 (Shannon et al., 2020; Touret et al., 2021, 2020; Weiss et al., 2021). The assay was performed in VeroE6 TMPRSS2 cells and calibrated in such a way that the cell culture supernatants were harvested (at 48 hours post infection) during the logarithmic growth phase of viral replication. The antibodies were tested in triplicate using 2-fold step-dilutions from 1000 to 0.97□ng/mL and from 5000 to 2.4 ng/mL for Cilgavimab and Tixagevimab alone and in combination. The amount of viral RNA in the supernatant medium was quantified by qRT-PCR to determine the 50% maximal effective concentration (EC_50_). Results were then compared with recent preliminary reports exploring the ability of the Omicron variant to escape neutralization by monoclonal antibodies.

We first observed a complete loss of detectable neutralizing activity for Casirivimab and Imdevimab (Roche-Regeneron), Bamlanivimab and Etesevimab (Eli-Lilly) and Regdanvimab (Celltrion) under our test conditions (Fig.1), which made it impossible to calculate EC_50_ (Table 1). This result is in line with previous EC_50_ determination reports (Aggarwal et al., 2021; Cameroni et al., 2021; Planas et al., 2021; VanBlargan et al., 2021; Xie et al., 2021) and with studies exploring the impact of amino-acid mutations in the SARS-CoV-2 spike receptor binding domain (RBD) conferring resistance to monoclonal antibodies (Dong et al., 2021; Starr et al., 2021a, 2021b).

**Table 1:** Interpolated EC_50_ values of therapeutic antibodies against SARS-CoV-2 BavPat1 and Omicron strains. EC_50_ values are expressed in ng/mL. For Sotrovimab, Cilgavimab and Tixagevimab the EC_50_ is the mean of two independent experiment (n=2), each including three replicates. (n.n: non-neutralising at highest concentration tested; n.t.: not tested, * antibodies were produced by the authors and are not the actual therapeutic products; # tested using a pseudovirus-based methodology).

|  |  | <u>This study</u> |  | <u>Aggarwal et al., 2021</u> | <u>Planas et al., 2021*</u> | <u>VanBlargen et al., 2021</u> | <u>Xie et al., 2021#</u> | <u>Cameroni et al., 2021#*</u> |
| --- | --- | --- | --- | --- | --- | --- | --- | --- |
| <u>Antibody Strain</u> |  | <u>BavPat B.1</u> | <u>Omicron</u> |  |  |  |  |  |
| <u>Regeneron</u> | <u>Casirivimab (REGN10933)</u> | **8** | n.n | n.n | n.n | n.n | n.n | n.n |
|  | <u>Imdevimab (REGN10987)</u> | **9** | n.n | n.n | n.n | n.n | n.n | n.n |
| <u>Lilly</u> | <u>Bamlanivimab (Ly-COV555)</u> | **13** | n.n | n.n | n.n | n.n | n.n | n.n |
|  | <u>Etesivimab (Ly-CoV16)</u> | **49** | n.n | n.t | n.n | n.n | n.n | n.n |
| <u>Celltrion</u> | <u>Regdanvimab (CT-P59)</u> | **13** | n.n | n.t | n.n | n.n | n.t | n.n |
| <u>GSK/ Vir</u> | <u>Sotrovimab (vir-7831)</u> | **89** | **276** | **1059** | **917** | **373** | **181** | **260** |
| <u>AstraZeneca</u> | <u>Cilgavimab (AZD1061)</u> | **93** | **1472** | nn | **1213** | n.t | n.n | **2772** |
|  | <u>Tixagevimab (AZD8895)</u> | **26** | n.n | **3490** | **8305** | n.t | **6860** | n.n |
|  | <u>Evusheld (AZD7442)</u> | **35** | **1488** | n.t | **773** | n.t | n.t | **418** |

Sotrovimab/Vir-7831 (GlaxoSmithKline and Vir Biotechnology) retains a neutralizing activity against the Omicron variant (Figure 1) with an EC_50_ shifting from 89 to 276 ng/ml, *i*.*e*. a fold change reduction of 3.1 (Table 1) in comparison with the ancestral B.1 strain. This result is in accordance with preliminary reports (Table 1) and with data from Vir Biotechnology using a pseudotype virus harboring all Omicron spike mutations (Cathcart et al., 2021). The fact that Sotrovimab retains significant activity against the Omicron variant can be related to the fact that this antibody, which was originally identified from a SARS-CoV-1 survivor and was found to also neutralize the SARS-CoV-2 virus, does not target the Receptor Binding Motif (RBM) but a deeper and highly conserved epitope of RBD (Pinto et al., 2020).

**Figure 1:**
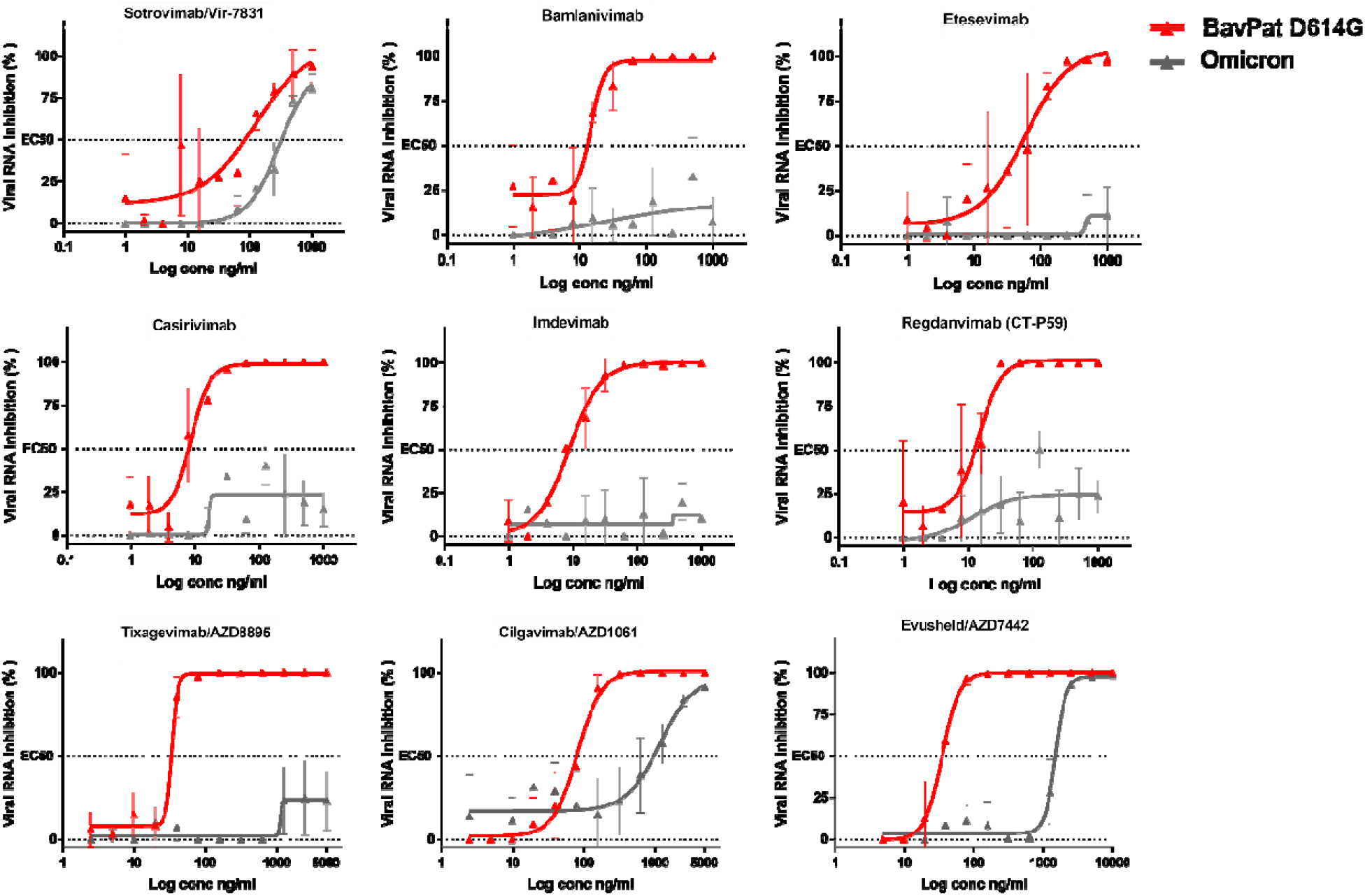
Dose response curves reporting the susceptibility of the SARS-CoV-2 BavPat1 D614G ancestral strain and Omicron variant to a panel of therapeutic monoclonal antibodies. Antibodies tested: Casirivimab/REGN10933, Imdevimab/REGN10987, Bamlanivimab/LY-CoV555, Etesevimab/LY-CoV016, Sotrovimab/Vir-7831, Regdanvimab/CT-P59, Tixagevimab/AZD8895, Cilgavimab/AZD1061 and Evusheld/AZD7742. Data presented are from three technical replicates in VeroE6-TMPRSS2 cells, and error bars show mean±s.d.

We found no significant neutralizing activity for Tixagevimab (EC_50_ >5000 ng/L) against Omicron as described in two other studies (Table 1). Cilgavimab conserved a neutralizing activity (Fig.1) with an EC_50_ shifting from 93 to 1472 ng/mL, *i*.*e*. a fold change reduction of 15.8, in accordance with Planas et al. (2021) (Table 1). When Cilgavimab was tested in combination with Tixagevimab, as proposed in the actual Evusheld/AZD7742 therapeutic cocktail (Mahase, 2021), the EC_50_ shifted from 35 to 1488 ng/mL, *i*.*e*. a fold change reduction of 42.6.

The observed decreases in activity should be seen in the context of the actual treatments given to patients. In the European Union, Sotrovimab is registered for the early treatment of infections (a single intravenous injection of 500 mg) and Evusheld is only registered at this stage for the prophylaxis of infection in subjects most at risk of developing severe forms of Covid-19 (150 mg Tixagevimab + 150 mg Cilgavimab, intramuscular). We defined a neutralization unit 50 (NU_50_), which is the amount of a given antibody needed to provide a 50% neutralization of 100 TCID_50_ of a given strain. We then calculated the number of neutralizing units present in each actual treatment proposed, based on the EC_50_s obtained previously, expressed in millions of neutralization units 50 per treatment (MNU_50_, Table 2).

**Table 2:** Neutralizing capacity of Sotrovimab, Cilgavimab and Evusheld. Values are expressed in millions of neutralizing units (MNU_50_) per treatment. One unit is defined as the amount of a given antibody needed to neutralize 50 % of 100 TCID_50_ of a given strain. Doses refer to treatments authorized in the European Union (Sotrovimab: 500 mg IV for the early treatment of infected patients (Gupta et al., 2021); Evusheld: 300 mg IM (corresponding to Cilgavimab 150 mg + Tixagevimab 150 mg) for prophylaxis of infection in patients with important risk factors).

|  | **BavPat B.1** | **Omicron** | **Fold change** |
| --- | --- | --- | --- |
| Sotrovimab/vir-7831 (500 mg) | **37.45** | **12.08** | □ **3.1** |
| Cilgavimab/AZD1061 (150 mg) | **10.75** | **0.68** | □ **15.8** |
| Tixagevimab/AZD8895 (150 mg) | **38.46** | n.n | - |
| Evusheld/AZD7442 (300 mg) | **57.14** | **1.34** | □ **42.6** |

The interest of this simulation is that it allows a realistic comparison of the neutralizing capacity of each treatment. Thus, the neutralizing capacity of a treatment with 300 mg of Evusheld against a type B.1 strain appears slightly higher than that conferred by 500 mg of Sotrovimab (57.14 *vs* 37.45 MNU_50_). In contrast, in the case of the Omicron variant, the neutralizing capacity of 300 mg Evusheld is about one tenth of that conferred by 500 mg Sotrovimab (1.3 *vs* 12.1 MNU_50_).

The activity of Evusheld against the BavPat1 B.1 European strain (57.14 MNU_50_) is slightly higher than that expected from the simple addition of the activities of Cilgavimab and Tixagevimab (10.75 and 38.46 MNU_50_, respectively, *i*.*e*. 49.21 MNU_50_) suggesting that if any synergistic action on different residues of the RBD exists, it is of modest magnitude. Against the Omicron strain, the activity of Evusheld (1.34 MNU_50_) is slightly higher than that of Cilgavimab alone (0.68 MNU_50_), which is consistent with the loss of a large part of the activity of Tixagevimab but may denote a limited complementation effect between the two antibodies. It remains therefore to be precisely documented by *in vivo* experiments whether the combination of Cilgavimab and Tixagevimab is preferable in clinical treatment to the use of Cilgavimab alone.

We conclude that, against the Omicron variant and compared to previous variants, Sotrovimab 500 mg retains a significant level of neutralizing activity. This activity is ∼30% of the activity of the same antibody treatment, and ∼20% of the activity of the Evusheld 300 mg cocktail, against a B.1 strain. The activity of Evusheld 300 mg against the Omicron variant is significantly reduced as it represents ∼10% of the activity of Sotrovimab 500 mg against Omicron, and ∼2.5% of the activity of the Evusheld cocktail against a B.1 strain. It will therefore be important to rapidly evaluate the actual therapeutic efficacy of Sotrovimab 500 mg and Evusheld 300 mg for the early treatment and prevention of infection with Omicron, respectively, at the doses initially proposed and to consider the possible need for dose modification or combination therapies.

## Materials & Methods

### Cell line

VeroE6/TMPRSS2 cells (ID 100978) were obtained from CFAR and were grown in minimal essential medium (Life Technologies) with 7⍰.5% heat-inactivated fetal calf serum (FCS; Life Technologies with 1% penicillin/streptomycin (PS, 5000U.mL^−1^ and 5000μg.mL^−1^ respectively; Life Technologies) and supplemented with 1⍰% non-essential amino acids (Life Technologies) and G-418 (Life Technologies), at 37°C with 5% CO_2_.

### Antibodies

Regdanvimab (CT-P59) was provided by Celltrion. Vir-7831 sotrovimab was provided by GSK (GlaxoSmithKline). The others antibodies: Bamlanivimab and Etesevimab (Eli Lilly and Company), Casirivimab and Imdevimab (Regeneron pharmaceuticals), Cilgavimab and Tixagevimab (AstraZeneca) were obtained from hospital pharmacy of the University hospital of La Timone (Marseille, France).

### Virus strain

SARS-CoV-2 strain BavPat1 was obtained from Pr. C. Drosten through EVA GLOBAL (https://www.european-virus-archive.com/) and contains the D614G mutation.

SARS-CoV-2 **Omicron** (B.1.1.529) was isolated from a nasopharyngeal swab of the 1^st^ of December in Marseille, France. Briefly, a 12.5cm^2^ culture flask of confluent VeroE6/TMPRSS2 cells was inoculated with the diluted sample. Cells were incubated at 37 °C during 6 hours after which the medium was changed with MEM medium with 5% FCS and incubation was continued for 3 days, until a CPE appeared. Supernatant was collected, clarified by spinning at 1500× g for 10 min, supplemented with 25mM HEPES (Sigma), aliquoted and stored at −80 °C. The full genome sequence has been deposited on GISAID: EPI_ISL_7899754 The strain, called 2021/FR/1514, is available through EVA GLOBAL (www.european-virus-archive.com, ref: 001V-04436)

All experiments with infectious virus were conducted in a biosafety level 3 laboratory.

### EC50 determination

One day prior to infection, 5×10^4^ VeroE6/TMPRSS2 cells per well were seeded in 100μL assay medium (containing 2.5% FCS) in 96 well culture plates. The next day, antibodies were diluted in PBS with ½ dilutions from 1000 to 0.97⍰ng/ml for most of them and from 5000 to 2.4 ng/ml for Cilgavimab and Tixagevimab. Eleven 2-fold or twelve 2-fold (for Cilgavimab and Tixagevimab) serial dilutions of antibodies in triplicate were added to the cells (25μL/well, in assay medium). Then, 25μL of a virus mix diluted in medium was added to the wells. The amount of virus working stock used was calibrated prior to the assay, based on a replication kinetics, so that the viral replication was still in the exponential growth phase for the readout as previously described (1–3). Four virus control wells were supplemented with 25μL of assay medium. Plates were first incubated 15 min at room temperature and then 2 days at 37°C prior to quantification of the viral genome by real-time RT-PCR. To do so, 100μL of viral supernatant was collected in S-Block (Qiagen) previously loaded with VXL lysis buffer containing proteinase K and RNA carrier. RNA extraction was performed using the Qiacube HT automate and the QIAamp 96 DNA kit HT following manufacturer instructions. Viral RNA was quantified by real-time RT-qPCR (GoTaq 1-step qRt-PCR, Promega) using 3.8μL of extracted RNA and 6.2μL of RT-qPCR mix and standard fast cycling parameters, *i*.*e*., 10min at 50°C, 2 min at 95°C, and 40 amplification cycles (95°C for 3 sec followed by 30 sec at 60°C). Quantification was provided by four 2 log serial dilutions of an appropriate T7-generated synthetic RNA standard of known quantities (10^2^ to 10^8^ copies/reaction). RT-qPCR reactions were performed on QuantStudio 12K Flex Real-Time PCR System (Applied Biosystems) and analyzed using QuantStudio 12K Flex Applied Biosystems software v1.2.3. Primers and probe sequences, which target SARS-CoV-2 N gene, were: Fw: GGCCGCAAATTGCACAAT; Rev: CCAATGCGCGACATTCC; Probe: FAM-CCCCCAGCGCTTCAGCGTTCT-BHQ1. Viral inhibition was calculated as follow: 100* (quantity mean VC-sample quantity)/ quantity mean VC. The 50% effective concentrations (EC50 compound concentration required to inhibit viral RNA replication by 50%) were determined using logarithmic interpolation after perorming a nonlinear regression (log(agonist) vs. response -- Variable slope (four parameters)) as previously described (2–6). All data obtained were analyzed using GraphPad Prism 7 software (Graphpad software).

## Contribution

FT and XDL conceived the experiments. XDL proposed the study. FT, CB, and HSB performed the experiments. FT, CB, HSB and XDL analyzed the results. FT and XDL wrote the paper. FT, CB, HSB and XDL reviewed and edited the paper.

## Acknowledgments

We thank Pr C Drosten for providing the SARS-CoV-2 BavPat strain through EVA GLOBAL. We thank the Noemie Courtin for the technical help regarding the antiviral assay and the sequencing.

